# Parallel Opposing GABAergic Pallidothalamic Pathways

**DOI:** 10.64898/2026.06.08.730902

**Authors:** Isaac Y.M. Chang, Jeanne T. Paz

## Abstract

Coordinated movement relies on interactions between the basal ganglia and the thalamus, yet the circuits linking these structures are not fully understood. Here, we dissect a non-canonical basal ganglia output pathway, from the external globus pallidus (GPe) to the nucleus reticularis thalami (nRT). We show that GPe neurons synapse onto all nRT neurons with 100% connectivity. Strikingly, despite its GABAergic nature, GPe input excites the posterior nRT but inhibits the anterior nRT, due to region-specific expression of the chloride transporter KCC2. This divergence shapes downstream thalamocortical activity, producing feedforward inhibition in the somatosensory thalamus but not the motor thalamus. Functionally, the in vivo activity of GPe→nRT axons is time-locked to movement. Overall, our work highlights the GPe as a major regulator of thalamic activity and reveals two parallel GPe→nRT streams with opposing effects on thalamic processing.

## Introduction

Movement is orchestrated by interconnected loops between the cortex and subcortical regions. The cortico-basal-ganglia-thalamo-cortical loop, in particular, is a fundamental circuit motif in vertebrates, and its anatomical connectivity is evolutionarily conserved across species(*1*, *2*). The rate model(*3*, *4*), formulated in the 1980s, provided a functional and structural framework for the corto-basal-ganglia-thalamo-cortical loop in motor function and dysfunction. A key component in this model is that basal ganglia output converges at the level of two nuclei, the internal globus pallidus (GPi) and the substantia nigra pars reticulata (SNr). The GPi/SNr, in turn, projects to the motor thalamus. In this working model, the GPi/SNr→motor thalamus pathway is thought to be the sole canonical basal ganglia output. Accordingly, one would predict that disruption in this canonical pathway would be detrimental to motor function. However, focal lesions or inactivation of the GPi performed on human Parkinson’s disease patients(*5*) or non-human primates(*6*) do not cause motor disturbances. This and other lines of evidence(*7*) challenge the notion that GPi/SNr are the sole basal ganglia output nuclei and raise the possibility of an additional circuit node that interfaces between the basal ganglia and the thalamus.

The projection pathway from the external globus pallidus (GPe) to the nucleus reticularis thalami (nRT) is of particular interest as a critical junction that bridges information transfer between the basal ganglia and the thalamus. The GPe is a central nucleus in the basal ganglia and sends widespread GABAergic projections to all basal ganglia nuclei, nRT, and regions beyond(*8*). The nRT sends powerful inhibitory connections to all thalamocortical (TC) nuclei, thereby gating information flow from the thalamus to the cortex(*9*). The GPe and nRT play central roles in regulating the network activity of the basal ganglia(*10*) and the thalamus(*11*), respectively. Via the GPe→nRT→TC pathway, the GPe is well-positioned to exert powerful control over TC activity. The existence of the GPe→nRT pathway was first shown in rats(*12–15*), cats(*16*), and squirrel monkeys(*17*, *18*) in the 1970s, and subsequent anatomical studies(*19–23*) followed. Dissecting the GPe→nRT pathway in isolation has been challenging due to the cellular heterogeneity of the GPe(*8*, *24*, *25*). Previous studies examining GPe regulation of nRT activity(*26–29*) or its behavioral role(*30*) largely relied on non-specific pharmacological, electrical, or optogenetic manipulations that engaged multiple GPe populations simultaneously. Given the numerous molecular subclasses of GPe neurons, each with distinct projection patterns, molecular identities, and electrophysiological properties(*8*, *25*), it has been difficult to parse out the specific contributions of distinct GPe pathways.

Here, we provide a comprehensive circuit characterization of the GPe→nRT pathway and uncover a mechanism by which GABAergic GPe input elicits region-specific postsynaptic responses of opposite polarity across the nRT, driven by local differences in chloride homeostasis.

## Results

### GPe sends widespread projections to the entire nRT

To visualize the GPe→nRT projection, we injected adeno-associated virus (AAV) carrying Cre-dependent eYFP into the GPe of Npas1-iCre mice **(Figure 1A)**. Npas1-positive (Npas1^+^) neurons comprise 32.1% of the total GPe population and label GPe→nRT projection neurons(*31*). We observed dense axonal arborizations distributed throughout the entire nRT, whereas TC nuclei were completely devoid of eYFP signal **(Figure 1B)**. Given the robust and selective labeling achieved with this mouse line, we used the Npas1-iCre mice for all subsequent experiments.

**Fig 1.**
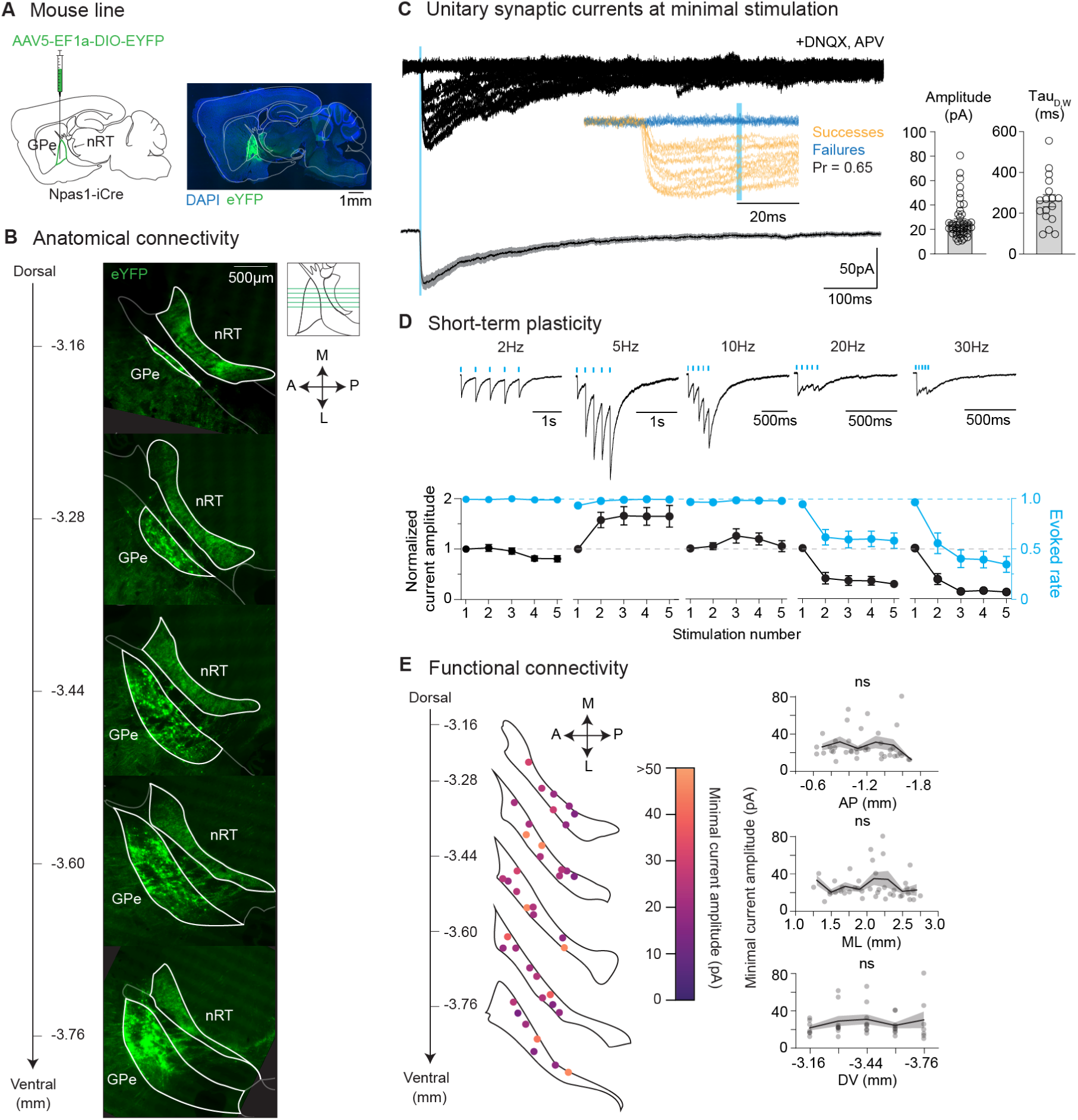
GPe sends widespread projections to the entire nRT. **A.** Injection schematics. **B.** Representative images of horizontal brain sections showing the presence of eYFP^+^ GPe axons throughout the entire nRT. Scale bar applies to all panels. **C.** Top, Representative traces of minimal oPSCs recorded from an nRT neuron in response to activation of a single GPe axon. Inset, expanded view of the traces. Stimulation intensity was adjusted to achieve a ∼50% success rate. Bottom, Mean trace ± SEM. Right, Electrophysiological features of minimal oPSCs: Amplitude (*n* = 47 cells) and weighted tau (*n* = 17 cells)**. D.** Top, Representative traces of oPSCs in an nRT neuron evoked by a train of light pulses delivered at 2 Hz (*n* = 20 cells), 5 Hz (*n* = 19 cells), 10 Hz (*n* = 19 cells), 20 Hz (*n* = 18 cells), and 30 Hz (*n* = 18 cells) at 1.5× the minimal stimulation intensity. Traces are scaled such that the amplitudes of the first evoked oPSC are the same. Bottom, Normalized current amplitudes and evoke rates across stimulation numbers. **E.** Left, Connectivity map showing the locations of recorded cells (*n* = 41 cells), and their corresponding amplitudes of minimal oPSCs. Right, Amplitudes of minimal oPSCs across the anterior-posterior (AP), medio-lateral (ML), and dorso-ventral (DV) axes. Note the uniformity of synaptic strength across all three axes. Linear regression (AP: *P* = 0.60; ML: *P* = 0.99; DV: *P* = 0.60).

To functionally characterize GPe→nRT synapses, we injected AAV carrying Cre-dependent ChR2-eYFP into the GPe of Npas1-iCre mice. Six weeks post-injection, we prepared acute horizontal brain slices and measured optically evoked postsynaptic currents (oPSCs) in nRT neurons **(Figures 1C and D)**. We recorded GABA-receptor mediated currents in nRT neurons in the presence of blockers of AMPA and NMDA receptors, DNQX (30µM) and APV (100µM), respectively. We optically activated GPe terminals in the nRT using the minimal stimulation intensity, defined as the light power that resulted in a 25–75% success rate. Synaptic currents recorded in this configuration are the result of the activation of a single synapse and reflect the quantal content(*32*). Paired-pulse stimulations revealed no correlation between the paired-pulse ratio and facilitation **(Figure S1)**, consistent with unitary synaptic transmission. oPSCs exhibit exceptionally slow decay kinetics with a tau decay constant of 263.61 ± 29.42 ms **(Figure 1C)**, typical of GABA_A_ currents in nRT neurons(*33*). Short-term plasticity of the GPe→nRT synapses was assessed by applying a train of five stimulations at frequencies ranging from 2 to 30Hz. GPe→nRT synapses show activity-dependent short-term facilitation, with the strongest synaptic facilitation at 5 Hz **(Figure 1D)**, which falls within the range of the *ex vivo* spontaneous firing rate of Npas1^+^ GPe neurons(*31*). We systematically recorded oPSCs across the entire nRT **(Figure 1E)**, and found, remarkably, that GPe stimulation evoked oPSCs in 100% (41/41) of recorded nRT neurons, irrespective of their location. Minimal oPSC amplitudes are uniformly distributed across all the anterior-posterior, medial-lateral, and dorso-ventral axes **(Figure 1E)**, consistent with GPe providing broad and spatially pervasive synaptic input throughout the nRT.

### Region-specific KCC2 expression drives opposing GABAergic GPe→nRT signaling

To assess how GPe input modulates nRT activity, we performed tight-sealed cell-attached recordings to monitor the firing changes of nRT neurons in response to optical stimulation of GPe→nRT terminals, without perturbing the intracellular milieu **(Figure 2A)**. GPe stimulation reduced firing in 21.3% (13/61) of nRT neurons, and did not affect firing in 26.2% (16/61) of nRT neurons. Surprisingly, activation of GABAergic GPe input increased firing in 52.5% (32/61) of nRT neurons **(Figure 2B)**. This finding was unexpected as GABAergic transmission is typically inhibitory in the adult brain, and examples of excitatory GABAergic synapses capable of directly driving firing are rare(*34–36*). Mapping the locations of recorded cells revealed a striking spatial segregation in GPe-evoked response: cells excited by GPe input were predominantly located in the lateral-posterior nRT, whereas those in the anterior-lateral nRT were more likely to be inhibited **(Figure 2D)**, revealing opposing effects of GPe input across these two regions.

**Fig 2.**
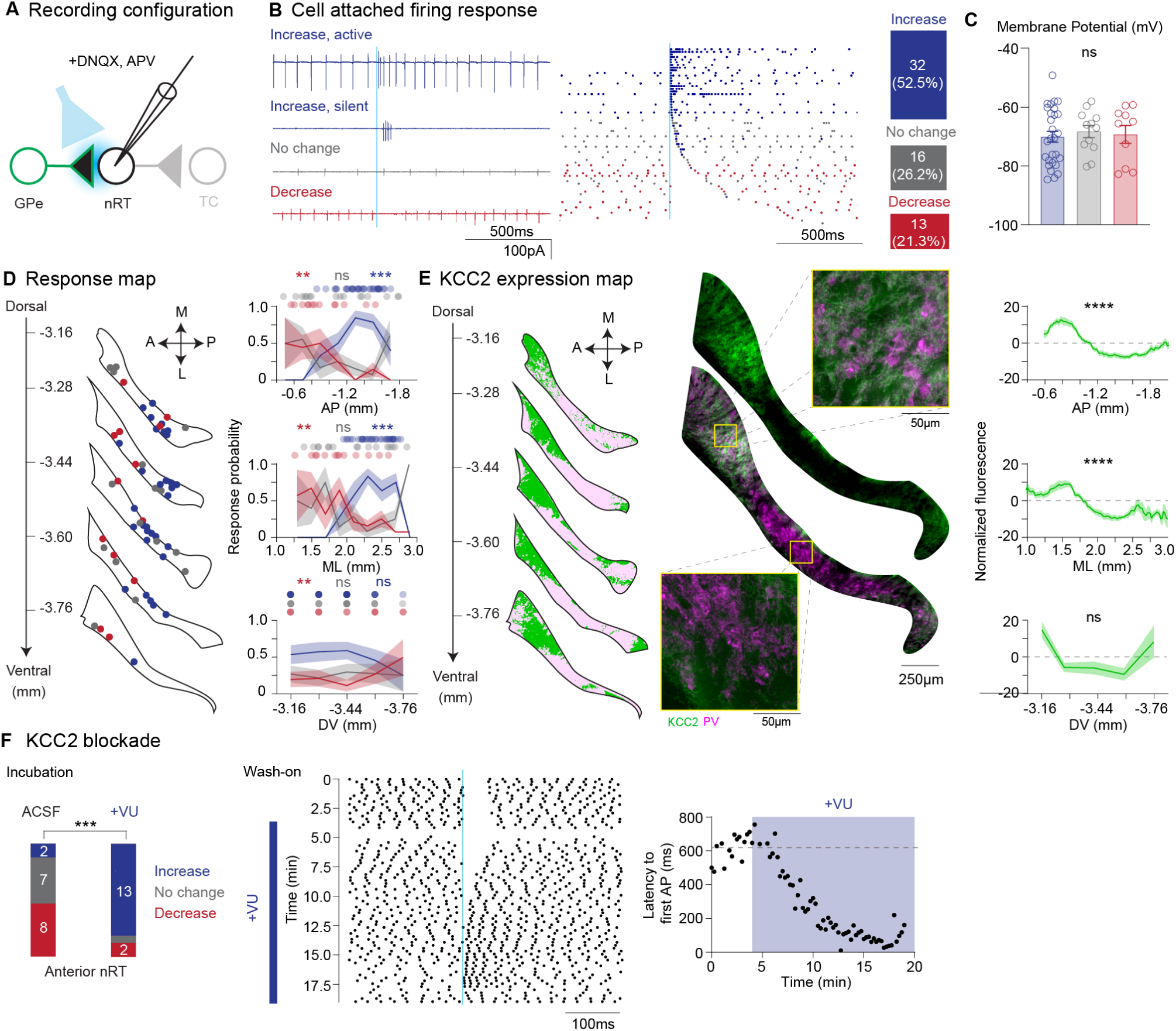
Region-specific KCC2 expression drives opposing GABAergic signaling from the GPe within the nRT. **A.** Recording configuration. **B.** Left, Representative traces of cell-attached recordings of nRT neurons in response to GPe axon stimulation. Middle, Raster plot showing spiking activity of all cells recorded, sorted based on the latency to the first spike after stimulation. Right, Numbers and percentages of response types. **C.** Resting membrane potential values across responders. Kruskal-Wallis test (*n* = 50 cells, *P* = 0.94). **D.** Left, Response map showing the locations of recorded cells (*n* = 61 cells) and their corresponding firing response. Right, Response probability across the AP, ML, and DV axes. Cells were predominantly inhibited in the anterior-medial regions and excited in the posterior-lateral regions of the nRT. Logistic regression (AP: Increase *P* = 1.34 × 10^−4^, No change *P* = 0.17, Decrease *P* = 2.68 × 10^−3^; ML: Increase *P* = 2.41 × 10^−4^, No change *P* = 0.18, Decrease *P* = 5.10 × 10^−3^; DV: Increase *P* = 0.39, No change *P* = 0.88, Decrease *P* = 7.44 × 10^−3^). **E**, Left, KCC2 expression map based on immunohistochemical signal from a representative animal. Middle, Representative images of horizontal brain sections. High-magnification image shows that KCC2 localizes to the perisomatic region and dendrites. Right, KCC2 fluorescence signal, normalized to the average signal of that section (*n* = 5 mice). KCC2 is restricted to the anterior nRT. Linear regression (AP: *P* = <1.00 × 10^−10^; ML: *P* = <1.00 × 10^−10^; DV: *P* = 0.50). **F.** Left, Number of response types with and without VU incubation in the anterior nRT. Fisher’s exact test (no VU vs. with VU: *n* = 33 cells, *P* = 1.91 × 10^−4^). Middle, Raster plot showing the effect of washing on VU 0463721 (KCC2 antagonist) on GPe-evoked response in an nRT neuron. Spiking was initially inhibited by GPe stimulation, and pharmacological blockade of KCC2 transformed GPe inhibition to excitation. Right, Effect of VU as quantified by the latency to the first spike after stimulation. Horizontal dashed line indicates the baseline latency.

How could a single source of GABAergic pallidal input produce this divergence in firing response? The effect of GABA_A_ receptor activation is determined by the difference between the membrane potential (Vm) and the chloride reversal potential (E_Cl_). To investigate whether a more hyperpolarized Vm at rest could be responsible for the increased firing (as GABA is depolarizing if and only if E_Cl_ > Vm), we measured Vm upon rupturing the membrane into whole-cell mode. Vm was similar between the three groups of responders **(Figure 2C)**, suggesting that the excitatory effects of GABA did not arise from differences in Vm.

Next, we tested the hypothesis that regional differences in chloride homeostasis, specifically mediated by KCC2, underlie the mixed responses upon GPe stimulation. KCC2 is a neuron-specific transporter that extrudes chloride ions (Cl⁻) from the cell, thereby maintaining a low intracellular chloride concentration, which is essential for hyperpolarizing GABAergic signaling in the adult brain. In rodents, it has been shown that the anterior and dorsal regions of the nRT are immunopositive for KCC2, but KCC2 is not present in other regions of the nRT(*37*, *38*).

We performed KCC2 immunostaining on mouse horizontal brain sections and found a clear anterior-posterior gradient of KCC2 expression within the nRT **(Figure 2E)**. KCC2 immunoreactivity was strong in the anterior nRT but was minimal in the posterior region. The expression pattern of KCC2 closely mirrored the GPe-evoked response pattern along the anterior-posterior axis **(Figure 2D)**. These data are consistent with the model in which high KCC2 expression in the anterior nRT maintains low intracellular chloride and a hyperpolarized E_Cl_, leading to inhibitory GABAergic responses, whereas reduced KCC2 expression in the posterior nRT leads to elevated intracellular chloride, depolarized E_Cl_, and excitatory GABAergic responses.

To test whether KCC2 determines the polarity of GPe-evoked responses, we pharmacologically blocked KCC2 using the selective antagonist VU 0463271 (100µM) and examined whether GPe-mediated inhibition could be converted into excitation under KCC2 blockade. We quantified the response profiles in anterior nRT from slices incubated with or without VU. Under control conditions, the majority of anterior nRT neurons were inhibited by GPe stimulation. KCC2 blockade increased the proportion of neurons exhibiting activation in response to GPe stimulation **(Figure 2F)**, suggesting that KCC2 activity constrains GABAergic signaling toward inhibitory responses at the population level. To determine whether this effect could occur within individual neurons, we performed VU wash-on experiments in anterior nRT neurons during steady spontaneous firing. Optogenetic stimulation of GPe axons initially suppressed spiking, but following the application of VU, the response progressively shifted from inhibition to excitation, and ultimately drove robust burst firing in the same cell **(Figure 2F)**. Together, these findings indicate that KCC2 activity is a key determinant of the polarity of GABAergic GPe input to nRT neurons.

### E_Cl_ governs the polarity of GPe-evoked firing response in nRT neurons

Does the observed KCC2 expression pattern translate to underlying differences in E_Cl_ in nRT neurons? We performed gramicidin-perforated patch recordings to measure physiological cell E_Cl_. Gramicidin is a cation-selective ionophore that preserves native intracellular chloride concentrations while permitting electrical access to the cell, thereby enabling accurate measurement of E_Cl_(*39*). We took advantage of the slow time course of perforation to first assess GPe-evoked firing response in cell-attached configuration following seal formation **(Figure 3A)**. This approach enabled direct correlation between firing response and the subsequently measured E_Cl_ within the same cell.

**Fig 3.**
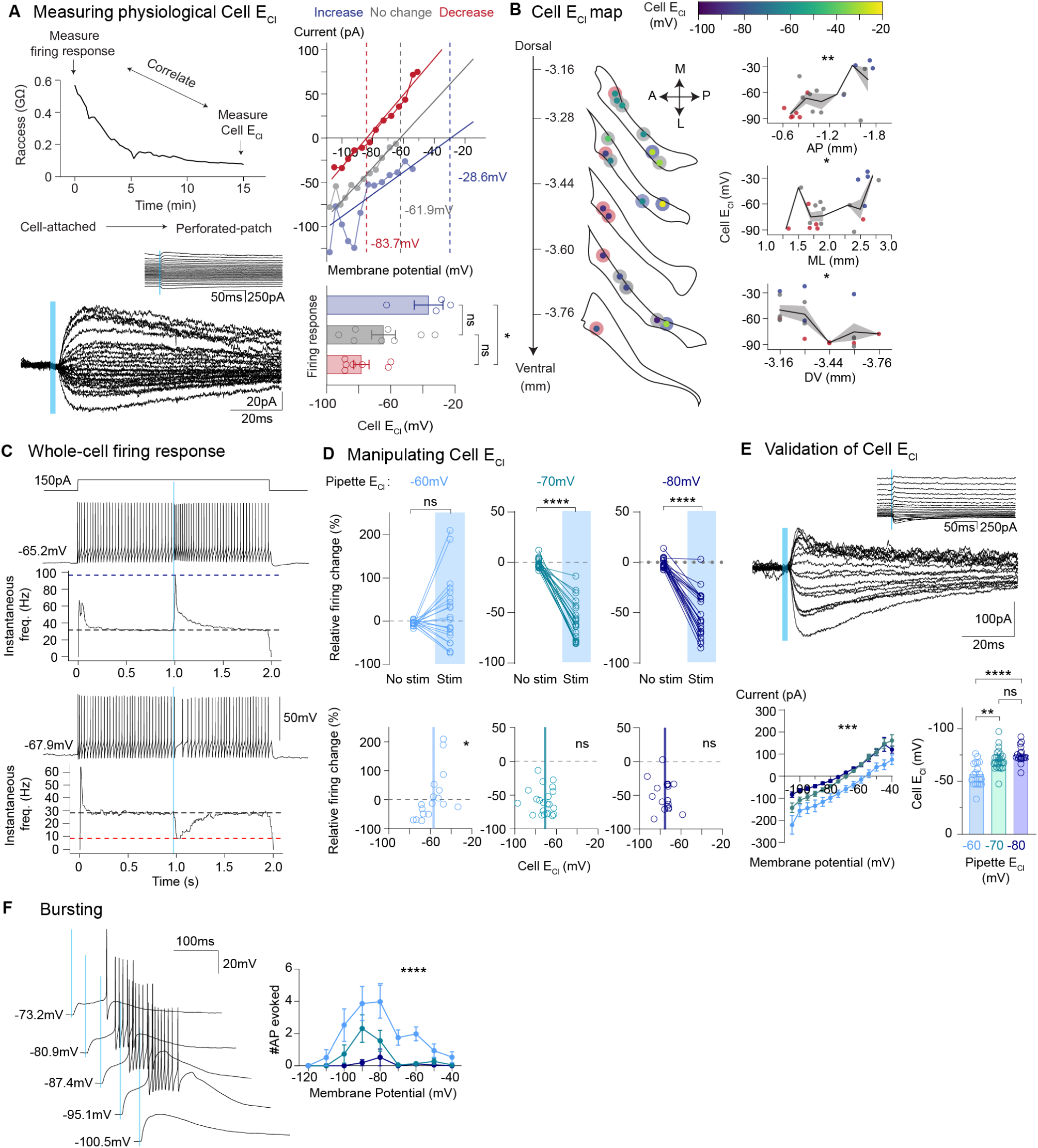
E_C_l governs the polarity of GPe-evoked response in nRT neurons. **A,** Top left, Recording protocol to assess physiological cell E_Cl_ and GPe-evoked firing response within the same cell. Access resistance was continuously monitored every 15 s after seal formation until desirable perforation was achieved. Bottom left, Representative traces of a gramicidin perforated-patch recording. Inset, same traces shown on a longer timescale and without baseline subtraction. Top right, Current–voltage relationship curves of three representative cells that showed a firing decrease (red), no change (gray), firing increase (blue) prior to E_Cl_ measurement. Vertical dashed lines indicate the x-intercepts. Numbers correspond to the measured cell E_Cl_ values. Bottom right, Cell E_Cl_ values across responders. Kruskal-Wallis test (*n* = 19 cells, *P* = 2.31× 10^−2^). Dunn’s multiple comparisons test (Increase vs. No change: *P* = 0.23; Increase vs. Decrease: *P* = 2.54 × 10^−2^; No change vs. Decrease: *P* = 0.83). **B,** Left, Cell E_Cl_ map showing the locations of recorded cells (*n* = 19 cells) and the correlation between cell E_Cl_ and firing response. Inner circles are color-coded to E_Cl_ values. Outer circles are color-coded to the firing response. Right, Cell E_Cl_ values across the AP, ML, and DV axes. Dots are color-coded to their firing response. Linear regression (AP: *P* = 4.42 × 10^−3^; ML: *P* = 2.10 × 10^−2^; DV: *P* = 2.21 × 10^−2^). **C.** Top, Representative trace of a whole-cell recording of an nRT neuron in response to GPe axonal stimulation during firing. The cell was recorded with an E_Cl_ = –60 mV pipette internal, which, upon GPe stimulation, increased in firing. Bottom, same as Top, but for a cell that was recorded with an E_Cl_ = –80 mV pipette internal that decreased firing upon stimulation. Black horizontal dashed line indicates the baseline firing rate. Blue or red horizontal dashed line indicates the poststimulus firing rate. **D,** Top, Firing responses of cells recorded with different internals: E_Cl_ = –60 mV, –70 mV, –80 mV. Wilcoxon test (–60 mV: *n* = 21 cells, *P* = 0.43; –70 mV: *n* = 23 cells, *P* = 2.38 × 10^−7^; –80 mV: *n* = 22 cells, *P* = 4.77 × 10^−7^). Bottom, Correlation between firing rate, shown in Top, and experimentally measured Cell E_Cl_ shown in E. Linear regression (–60 mV: *n* = 17 cells, *P* = 1.61 × 10^−2^; –70 mV: *n* = 22 cells, *P* = 0.80; –80 mV: *n* = 19 cells, *P* = 0.77). **E,** Top, Representative traces of oPSCs recorded at different command potentials. Inset, same traces shown on a longer timescale and without baseline subtraction. Bottom left, I-V curves. Mixed ANOVA (*n* = 60 cells, *P* = 5.54 × 10^−4^). Bottom right, Experimentally measured cell E_Cl_ values, which closely match their respective pipette E_Cl_ values. Kruskal-Wallis test (*n* = 64 cells, *P* = 1.15 × 10^−5^). Dunn’s multiple comparisons test (–60 mV vs. –70 mV: *P* = 2.23 × 10^−3^; –60 mV vs. –80 mV: *P* = 9.62 × 10^−6^; –70 mV vs. –80 mV: *P* = 0.31). **F,** Left, Representative traces showing evoked burst firing in an nRT neuron in response to GPe stimulation. Right, Number of action potentials evoked across different membrane potentials in the three pipette E_Cl_ conditions. RM two-way ANOVA (*n* = 68 cells, *P* = 2.68 × 10^−5^).

After characterizing GPe-evoked response in cell-attached configuration, we monitored access resistance until stable perforation was achieved. E_Cl_ was determined from GPe-evoked currents recorded across command potentials **(Figure 3A)**. Current-voltage (I-V) relationship curves were constructed, and x-intercepts of linear fits were used to determine E_Cl_. Cells excited by GPe stimulation exhibited significantly more depolarized E_Cl_ than inhibited cells **(Figure 3A)**. Anatomical mapping again revealed a pronounced anterior-posterior gradient in E_Cl_ **(Figure 3B)**, closely mirroring the KCC2 expression pattern **(Figure 2E)** and the distribution of excitatory versus inhibitory responses **(Figure 2B)**.

We next tested whether we could control the directionality of nRT neurons’ response to GPe by varying intracellular chloride in whole-cell configuration. Whole-cell recordings were performed using pipette internal solutions of different E_Cl_: –60 mV, –70 mV, and –80 mV. Experimental measurement of cell E_Cl_ obtained from I-V curves closely matched the predicted pipette E_Cl_ **(Figure 3E)**. We injected +150 pA current to depolarize the cell to an active state and measured the relative change in firing rates upon GPe stimulation **(Figure 3C and D)**. Using the –60 mV E_Cl_ internal, 52.4% (11/21) of the cells increased firing, consistent with the proportion of excited cells observed in cell-attached recordings **(Figure 2B)**. In contrast, using the –70 mV and –80 mV E_Cl_ internals, GPe stimulation consistently resulted in inhibitory postsynaptic potentials (IPSPs) and suppressed firing in all nRT cells **(Figure 3D)**. Together, these findings demonstrate that GABA release from GPe axons can exert either activating or suppressing effects depending on the E_Cl_ of nRT neurons.

Finally, we assessed the effects of GPe input on nRT neurons at rest, in the absence of ongoing firing. Consistent with our cell-attached recordings **(Figure 2B)**, GPe stimulation evoked high-frequency burst firing in the posterior nRT neurons **(Figure 3F)**. Notably, evoked action potentials ride on a slow and prolonged depolarizing current, presumably mediated by low-threshold Ca^2+^ channels (e.g., CaV3.3) expressed in nRT neurons(*40*). The number of action potentials increased with more depolarized E_Cl_ internals **(Figure 3F)**. The E_Cl_ values of all internals used were more hyperpolarized than the activation threshold of Na^+^ channels, suggesting that depolarizing GABA_A_ current first recruits low-threshold Ca^2+^ channels, which in turn trigger Na^+^-mediated action potentials. This is consistent with the finding that pharmacological blockade of CaV3.3 eliminates burst firing evoked by GABA puff in nRT neurons(*41*).

### GPe differentially regulates downstream thalamocortical activity through feedforward inhibition

Given the strong and regionally specialized control of GPe input on nRT activity, we next investigated whether GPe stimulation can differentially modulate the activity of TC neurons downstream of nRT **(Figure 4A)**. We recorded responses in two TC nuclei neighboring the nRT, the somatosensory ventrobasal nucleus (VB), which receives inputs from the posterior nRT, and the motor ventrolateral nucleus (VL), which receives inputs from the anterior nRT(*42*). GPe stimulation robustly inhibited firing in VB neurons and evoked burst-like IPSPs and inhibitory postsynaptic currents (IPSCs) in VB neurons **(Figure 4B)**, consistent with our finding that GPe stimulation evoked burst firing in the nRT neurons in the posterior region **(Figure 2B and 3F)**, which is known to project to the VB thalamus(*43*). In contrast, GPe stimulation produced little to no effect on the firing of VL neurons **(Figure 4C)**. Because nRT neurons are largely silent in acute slice preparations, GPe inhibition of the anterior nRT neurons **(Figure 2B)** would not substantially affect downstream VL activity.

**Fig 4.**
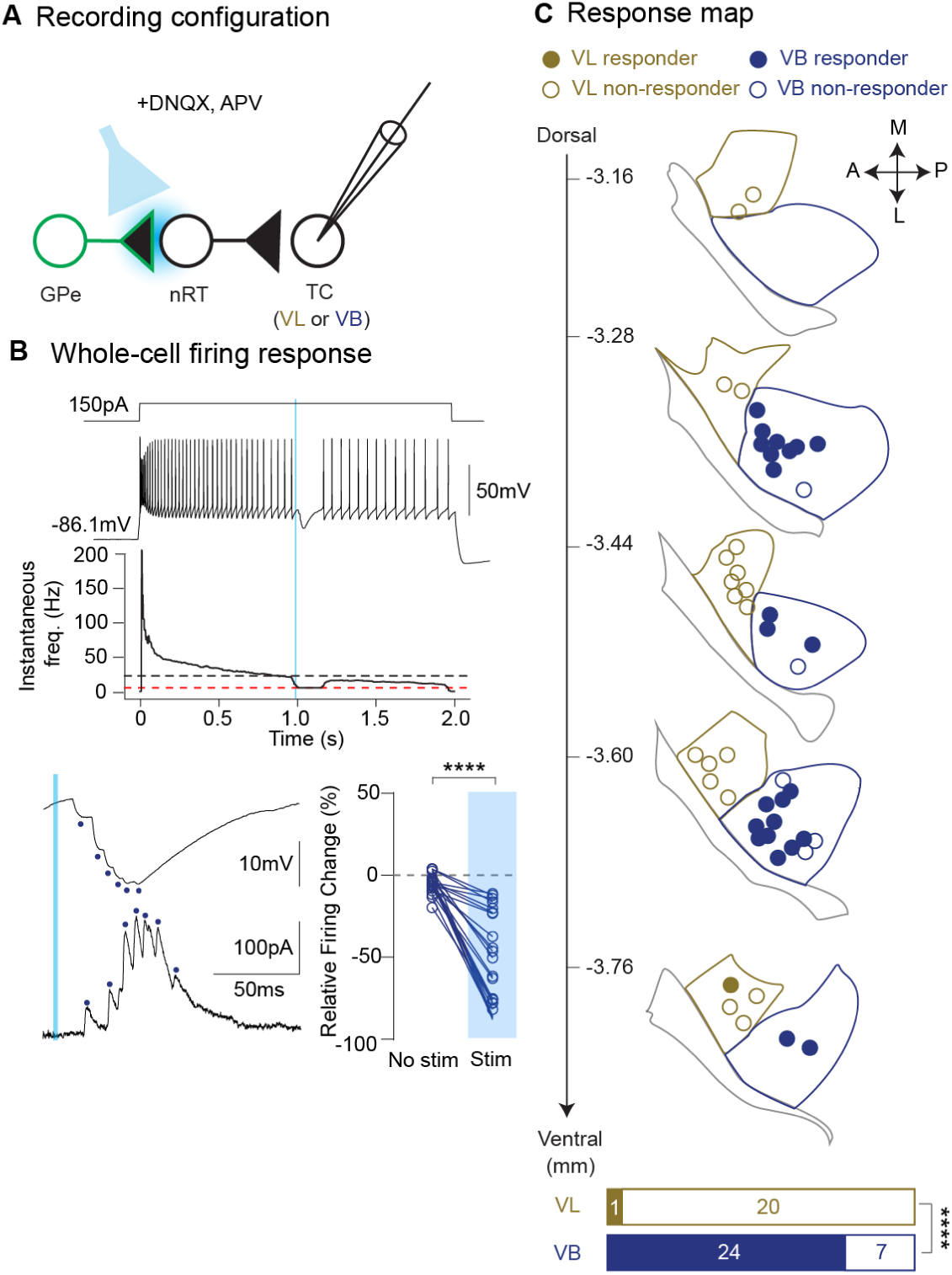
GPe differentially regulates downstream thalamocortical activity through feedforward inhibition. **A,** Recording configuration. **B,** Top, Representative trace of a whole-cell recording of a VB neuron in response to GPe axonal stimulation during firing. The cell was recorded with an E_Cl_ = –80 mV pipette internal, which, upon GPe stimulation, decreased in firing. Black horizontal dashed line indicates the baseline firing rate. Red horizontal dashed line indicates the poststimulus firing rate. Bottom left, expanded view of the traces showing evoked burst IPSPs. Blue dots denote the timing of individual IPSPs. In a separate cell, GPe evoked similar burst IPSCs in voltage clamp. Bottom right, Firing response. Wilcoxon test (*n* = 24 cells, *P* = 1.19 × 10^−7^). **C,** Response map showing the locations of recorded cells (*n* = 64 cells). Virtually no responders were observed in the VL nucleus. Fisher’s exact test (VL vs. VB: *P* = 2.67 × 10^−7^).

### *In vivo* activity of GPe→anterior nRT pathway tracks movement states and transitions

Given the central role of the GPe in basal ganglia circuitry and motor control, we asked whether the GPe→nRT pathway might contribute to the regulation of movement. Previous studies have shown phasic changes in GPe firing(*44–51*) and nRT firing(*52–55*) during movement. However, whether such activity extends to the Npas1⁺ subpopulation or specifically to GPe→nRT projection neurons remains unclear. Because the anterior nRT projects to the motor VL thalamus(*42*), we hypothesized that motor-related effects are mediated primarily by the GPe→anterior nRT→VL pathway. Accordingly, we focused on this pathway for our *in vivo* experiments.

To investigate the contribution of the GPe→anterior nRT pathway to motor behavior, we monitored the real-time activity of GPe axons in the anterior nRT using fiber photometry. We used an axon-targeted GCaMP variant, axon-GCaMP6s(*56*), to image axonal Ca^2+^ dynamics at GPe→anterior nRT terminals. Axonal Ca^2+^ signals serve as a proxy for presynaptic activity and neurotransmitter release(*57*), providing a pathway-specific readout of presynaptic output. Npas1-iCre mice were unilaterally injected with AAV carrying Cre-dependent axon-GCaMP6s into the GPe and implanted with an optical fiber in the anterior nRT **(Figure 5A)**. We monitored mouse speed during self-paced locomotion in an open-field arena **(Figure S2A)** and aligned GPe axonal Ca^2+^ activity traces to mouse speed **(Figure 5B)**. Activity in the GPe→anterior nRT pathway, quantified by the z-scored GCaMP fluorescence signal, increased at movement onset and decreased at movement offset **(Figure 5C)**. Analysis of the relationship between speed and GCaMP activity revealed a strong linear relationship at low movement speeds (0–1 cm/s), followed by a plateau at higher speeds **(Figure 5D)**. In this low-speed window, pathway activity correlated with speed, with a maximal Pearson correlation of 0.34 ± 0.04. The plateau in signal at higher speeds may reflect sensor saturation or a ceiling effect in GPe→nRT pathway recruitment.

**Fig 5.**
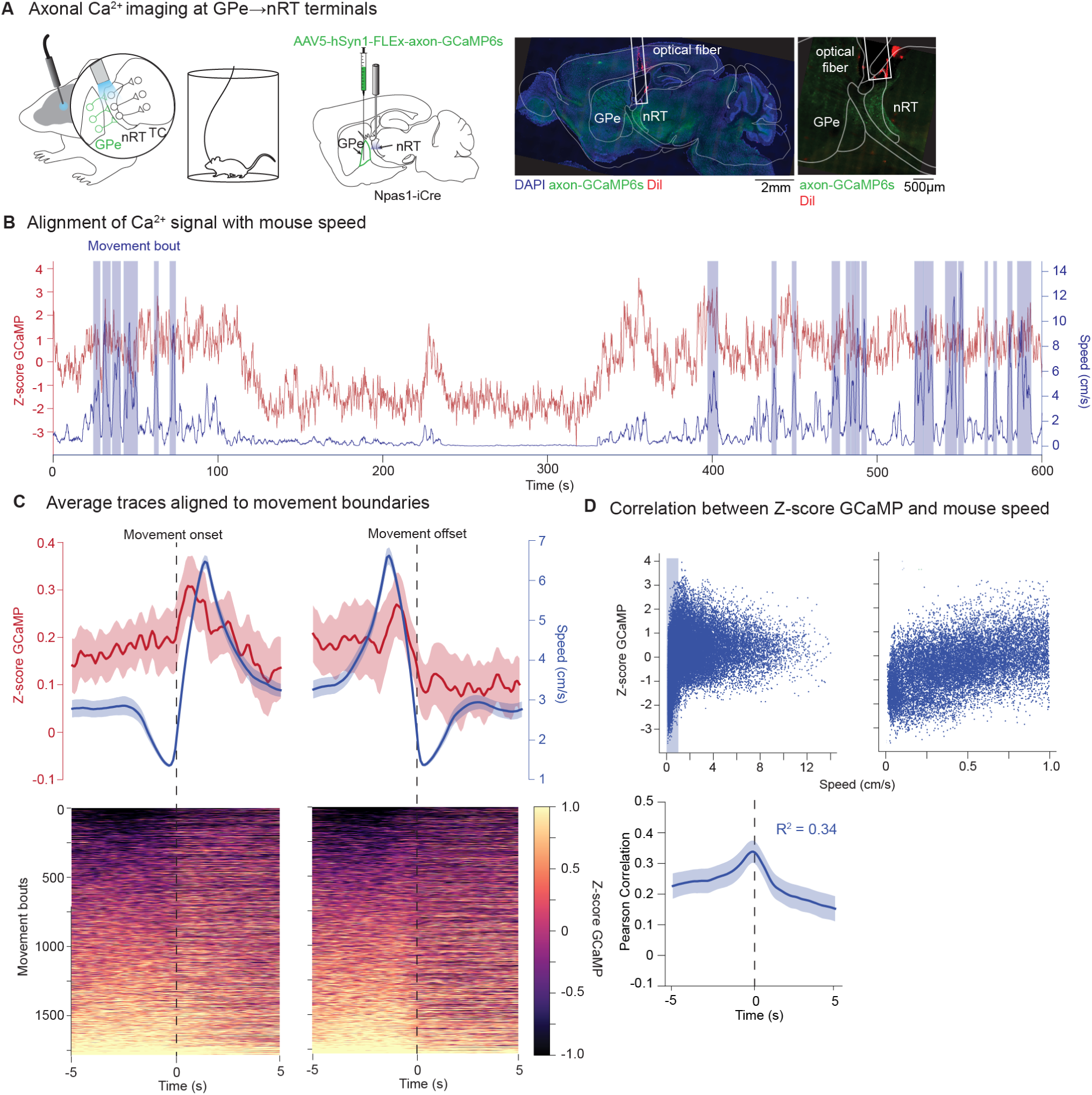
In vivo activity of GPe→anterior nRT pathway tracks movement states and transitions. **A,** Experimental design. **B,** Representative trace of mouse speed and Z-score GCaMP over time. Blue boxes indicate individual movement bouts. **C,** Z-score GCaMP aligned with movement onset (left, *n* = 1812 bouts, *N* = 10 mice) and offset (right, *n* = 1811 bouts, *N* = 10 mice). Top, Mean traces ± SEM. Bottom, Heatmaps of Z-score GCaMP sorted based on pre-onset/offset activity. **D,** Top, Scatter plot showing the relationship between Z-score GCaMP and mouse speed from one recording. Each marker is one frame (1/10 s). Blue rectangle indicates the speed window (0–1 cm/s) expanded on the right. Bottom, Time-resolved correlation between Z-score GCaMP and mouse speed for the chosen speed window. Mean traces ± SEM.

Because recordings were performed unilaterally from the right hemisphere, we also asked whether pathway activity was associated with directional turning behavior. We quantified angular velocity based on changes in the mouse heading direction **(Figure S2B)**. Both left and right turns were associated with transient increases in GCaMP activity, but no left/right bias was observed **(Figure S2C)**. Together, these results provide evidence that the GPe→anterior nRT pathway is dynamically coupled to ongoing movement states and transitions in freely behaving mice.

## Discussion

This study provides the first detailed analysis of the GPe→nRT pathway since its initial identification in the 1970s. We show that this non-canonical pallidothalamic pathway possesses several unique properties supporting strong modulation of thalamic activity and motor output: 1) widespread connectivity across the entire nRT and 2) region-specific GABAergic signalling that depends on an anterior-posterior expression of the chloride transporter KCC2. These properties position the GPe→nRT pathway as a critical circuit node in basal ganglia-thalamus communication.

### Opponent control of thalamic activity through selective activation and inhibition

One of the most unexpected findings is that the broad and near-uniform functional connectivity of GPe→nRT input **(Figure 1E)** did not translate into a homogeneous response across nRT regions: Anterior nRT neurons were inhibited, whereas posterior nRT neurons were excited in response to GPe stimulation **(Figure 2B)**. This bidirectional modulation of nRT activity was shaped by an anterior-posterior gradient in KCC2 expression, with high levels in the anterior nRT and low levels in the posterior nRT **(Figure 2E)**, consistent with a similar anterior-posterior gradient of E_Cl_ **(Figure 3B)**. Previous studies have reported a wide range of E_Cl_ in the nRT (*58*, *41*, *38*, *59*), ranging from −71.0 mV(*58*) to −45.0 mV(*41*). We show that the KCC2 expression gradient described here **(Figure 2E)** provides a mechanistic basis explanation for this variability of E_Cl_, and that E_Cl_ is spatially organized along the same anterior-posterior gradient as KCC2 expression **(Figure 3B)**.

What does the dual and opposing effect of GPe GABAergic input on the nRT mean for information gating in the thalamus? The nRT has long been proposed to function as a “searchlight” that selectively gates information flow through the thalamus, thereby shaping sensory processing, attention, and behavior(*11*). The original formulation proposes that this searchlight function is mediated through intrinsic intra-nRT connections, whereby localized nRT activity patterns selectively gate downstream TC nuclei. Our findings suggest an additional, complementary mechanism in which this searchlight function could be perhaps achieved through region-specific GABAergic excitation and inhibition from the GPe. Specifically, the inhibitory GPe→anterior nRT pathway and excitatory GPe→posterior nRT pathway may constitute parallel information streams that are recruited under certain behavioral conditions. This unique circuit architecture could enable flexible and context-dependent modulation of thalamic output(*60*).

In addition, the organization of the extrinsic GPe→nRT input differs from that of intrinsic intra-nRT connectivity. Whereas intra-nRT connections are relatively sparse, GPe→nRT input is pervasive and distributed throughout the nRT(*61*), suggesting these systems likely serve complementary roles. Intrinsic intra-nRT connectivity is well-positioned to mediate local interactions between neighboring nRT neurons and coordinate activity within regions. In contrast, the widespread nature of GPe→nRT suggests thalamic gating on a broader scale, dynamically regulating thalamic activity across regions.

One open question is the functional role of GPe input to the rest of the nRT, such as the posterior region, which is excited by GPe input and thus provides feedforward inhibition onto the VB thalamus **(Figure 4)**. Given the established role of VB in sensory processing(*43*, *62*), this pathway may contribute to sensory filtering mechanisms that suppress irrelevant or distracting inputs while enhancing the salience of behaviorally relevant signals. Consistent with this, broad optogenetic suppression of the GPe has previously been shown to enhance behavioral performance in an auditory discrimination task through the auditory nRT(*30*). The GPe→posterior nRT→VB pathway may similarly contribute to filtering background noise activity through feedforward inhibition and improve neural signal-to-noise ratio. Although the present study primarily focuses on motor-related activity of the GPe→anterior nRT pathway, defining the role of GPe→posterior nRT pathway will be an important future direction.

### Implications for basal ganglia-thalamus connectivity and function

The classical circuit model posits the GPi and the SNr as the canonical output nuclei of the basal ganglia, which give rise to GABAergic projections that inhibit the motor thalamus, thus inhibiting movement. Due to their characteristically high tonic firing rates *in vivo*(*63*, *64*), GPi/SNr are thought to tonically inhibit the motor thalamus(*65*). Movement initiation requires a transient pause in GPi/SNr firing, thereby disinhibiting the motor thalamus and allowing motor programs to be executed(*66*).

How the non-canonical GPe→nRT pathway operates alongside the canonical GPi/SNr pathway to coordinate movement remains to be determined. The activity of GPi/SNr is controlled upstream through a population of GPe neurons that are parvalbumin-positive and distinct from the Npas1^+^ GPe neurons that project to the nRT. The relative *in vivo* firing patterns of these two GPe subtypes have been studied in dopamine-depleted animals(*10*, *67*), but not under physiological conditions. Thus, it is currently difficult to determine whether these pathways operate in concert, in opposition, or in parallel. Simultaneous monitoring of both pathways during behavior will be essential to elucidate how they shape thalamic activity and motor output.

### Conclusions and outlook

This work expands the classical basal ganglia model by positioning the GPe as a direct regulator of thalamic activity alongside the canonical GPi/SNr output pathway. By establishing this novel route, our findings demonstrate that basal ganglia-thalamic communication is not a static inhibitory gate, but a dynamic system. These findings establish the GPe→nRT pathway as a previously underappreciated component of basal ganglia-thalamic circuitry with broad implications for thalamic function in health and disease.

## Methods

### Mice

We performed all experiments per protocols approved by the Institutional Animal Care and Use Committee at the University of California, San Francisco, and Gladstone Institutes. Precautions were taken to minimize stress and the number of animals used in all experiments. Adult (P56–P180) male and female mice were used. C57BL/6J mice (wild-type, IMSR_JAX:000664), PV-Cre mice (PV-Cre, IMSR_JAX:017320), SOM-Cre mice (SOM-IRES-Cre, IMSR_JAX:013044), Npas1-iCre (Npas1-Cre-2A-tdTomato BAC transgenic line 1, IMSR_JAX:027718, generously provided by Dr. Aryn Gittis, Carnegie Mellon University) were used.

### Viral injections and optic fiber implantations

Mice aged postnatal day 50–70 were anesthetized with 2–5% isoflurane and secured in a stereotaxic apparatus (Kopf Instruments). AAV constructs were delivered using the following coordinates into the GPe: (relative to bregma, AP: –0.28 mm; ML: 2.10 mm; DV: 4.00 mm, 150 nL, 30 nL/min); into the anterior nRT: (relative to bregma, AP: –0.70 mm; ML: 1.30 mm; DV: 3.70 mm and 4.10 mm at two DV positions, 40 nL each, 20 nL/min). For all injections, the MicroSyringe Pump Controller (Micro4, WPI), NanoFil syringes (10 μL, WPI), and 33-g beveled NanoFil needles (NF33BV-2, WPI) were used. After injection, the needle was left at the injection site for 5 min before slow withdrawal to allow diffusion and minimize backflow. Viral injections were performed unilaterally for *ex vivo* slice electrophysiology, circuit mapping studies, and fiber photometry. Experiments were performed at least six weeks after viral injections.

For circuit mapping experiments, we injected AAV5-EF1a-DIO-EYFP (RRID: Addgene_27056, 9.6 × 10^12^ GC/mL) into the GPe or nRT. For fiber photometry experiments, four weeks after viral injections, fiber optic cannulae (Doric Lenses, 1.25 mm zirconia ferrule, 400 µm core, 0.66 NA, 4.0 mm length) were implanted unilaterally using the same coordinates above the nRT. Cannulae were fixed to the skull using dental cement (Henry Schlein, Flow It ALC Flowable Composite A0). Optical fibers were coated with Dil stain to visualize fiber tracts.

### *Ex vivo* patch-clamp recordings

Mice aged postnatal day 90–140 (at least 6 weeks after viral injections) were anesthetized with 4% isoflurane and perfused transcardially with ice-cold sucrose slicing solution containing 234 mM sucrose, 26 mM NaHCO_3_, 11 mM glucose, 10 mM MgSO_4_, 2.5 mM KCl, 1.25 mM NaH_2_PO_4_, and 0.5 mM CaCl_2_, equilibrated with 95% O_2_ and 5% CO_2_. Brains were rapidly extracted and dissected in ice-cold sucrose slicing solution. Horizontal slices containing the thalamus were cut at a thickness of 250µm. We incubated the slices, initially at 32°C for 30 min and then at room temperature (24–26 °C) for 1 h, in artificial cerebrospinal fluid (ACSF) containing 126 mM NaCl, 26 mM NaHCO_3_, 10 mM glucose, 2.5 mM KCl, 2 mM CaCl_2_, 1.25 mM NaH_2_PO_4_, and 1 mM MgSO_4_, equilibrated with 95% O_2_ and 5% CO_2_.

Following incubation, brain slices were transferred to the recording chamber and superfused with ACSF at a flow rate of 2 mL/min at room temperature. We obtained recordings from nRT and TC neurons visually identified using differential contrast optics with an Olympus microscope (60X objective; numerical aperture, 1.1; working distance 1.5 mm; SKU 1-U2M592). Recording electrodes made of borosilicate glass had a resistance of 3–6 MΩ when filled with an internal solution. In all recording conditions, we monitored access resistance (Raccess), and cells were included for analysis only if the Raccess was <25 MΩ. All recordings were performed in the presence of DNQX (30 µM, Sigma-Aldrich, no. D0540) and DL-2-Amino-5-phosphonopentanoic acid (APV) (100 µM, Sigma-Aldrich, no. A5282) to pharmacologically isolate GABA_A_-mediated events. Electrophysiological data were acquired at 100 kHz and filtered at 3 kHz using a Multiclamp 700B amplifier (Molecular Devices) and the pClamp11 software suite (Molecular Devices).

For voltage-clamp recordings, the internal solution contained 135 mM CsCl, 10 mM HEPES, 10 mM EGTA, 5 mM QX-314 (lidocaine N-ethyl bromide), and 2 mM MgCl_2_, pH adjusted to 7.3 with CsOH (290 mOsm/kg). E_Cl_ was estimated to be about 0 mV based on the Nernst equation. Neurons were clamped at –65 mV. Junction potential, calculated using the JPCal Program in Clampex, was –3.2 mV and left uncorrected.

For cell-attached and current-clamp recordings, the internal solution contained 120 mM potassium gluconate, 11 mM KCl, 10 mM HEPES, 1 mM EGTA, 1 mM MgCl_2_, 1 mM CaCl_2_, pH adjusted to 7.4 with KOH (290 mOsm/kg). E_Cl_ was estimated to be about –60 mV based on the Nernst equation. Some experiments used internal solutions with –70 mV and –80 mV E_Cl,_ which contained the same ingredients but with 127 mM potassium gluconate/4 mM KCl, 129 mM potassium gluconate/2 mM KCl, respectively. We clamped the neurons at –80 mV for cell-attached recordings. The three internal solutions used had junction potential of –12.5 mV (E_Cl_: –60 mV), –13.2 mV (E_Cl_: –70 mV), and –13.7 mV (E_Cl_: –80 mV). Junction potentials were corrected offline. In a subset of experiments, KCC2 was blocked by bath application of the selective antagonist, VU 0463271 (100 µM, Sigma-Aldrich, SML2333).

For perforated-patch recordings, the internal solution contained 130 mM KCl, 10 mM HEPES, 10 mM EGTA, 1 mM MgCl_2_, 1 mM CaCl_2_, pH adjusted to 7.3 with KOH (290 mOsm/kg). The internal solution was first warmed to 37°C, and combined with a stock solution of gramicidin A (dissolved in DMSO, Sigma-Aldrich, no. 50845) to achieve a final concentration of 100µg/mL. The solution was then vortexed and sonicated for 1 min to ensure complete dissolution of gramicidin. Patch pipettes were prepared first by tip-filling with a gramicidin-free solution, and then back-filled with the gramicidin solution. To minimize gramicidin precipitation, the internal solution was vortexed every hour during experiments. Patch pipettes with a blunt taper were used to facilitate diffusion of gramicidin to the tip. After initial seal formation, perforation was then monitored by tracking Raccess, which was determined from the current response to a −10mV voltage drop. Mean Raccess at the start of E_Cl_ measurement was 71.28 ± 3.30 MΩ. Recordings that failed to reach these Raccess values within the first 10 minutes rarely achieved sufficient quality. Rupture or breakthrough of the patch was determined by a supraphysiological cell E_Cl_. Pipette E_Cl_ was estimated to be about 0 mV based on the Nernst equation, which is not physiologically plausible. Junction potential was –2.8 mV and left uncorrected. To account for voltage errors introduced by large Raccess, offline correction was performed. The voltage drop across the Raccess was calculated by multiplying the measured current response by the Raccess, and this value was subtracted from the command voltage to estimate the actual membrane potential.

### Fiber photometry

Fiber photometry was performed two weeks after optical fiber implantations. Behavioral tests were performed in a standard lit room between 10:00 am and 7:00 pm. Mice were tested in a white plastic open-field cylinder (diameter 30 cm), and was cleaned with 70% ethanol between sessions. On the day before the experimental session, each mouse was allowed to acclimate to the open field for 20 min, with a low-autofluorescence fiber optic cable attached to the implant via a rotary joint (Doric Lenses).

Recordings were performed on an RZ10X processor (Tucker-Davis Technologies) with 405, 465 LEDs. LED drivers were calibrated such that light power was approximately 15 μW for 405 nm and 20 µW for 465 nm at the tip of the optical fiber. For axon-GCaMP6s excitation, 405 nm and 465 nm LEDs were sinusoidally modulated at distinct frequencies (210 Hz and 330 Hz, respectively). Fluorescence signals and mouse behavior were recorded for 1 h. An overhead camera (Allied Vision) was used to acquire mouse behavior in the open field at 10 fps, 640 x 480 px per frame. Mouse position was tracked using Deeplabcut **(see Methods)**. After the completion of behavioral experiments, transgene expression and optical fiber locations were histologically verified *post hoc*. Only histologically validated subjects were included in this study. Mice with baseline locomotor activity less than 1 cm/s were excluded, as this is indicative of abnormal motor behavior, possibly caused by injury during surgical procedures.

### Markerless pose estimation

Behavioral videos were analyzed using DeepLabCut(*68*) (DLC) for markerless tracking of mouse body parts. DLC version 3.0.0rc9 was used with default parameters and a pretrained ResNet50. Seven body parts (nose, left ear, right ear, body center, left lateral, right lateral, tail base) were manually labelled. A total of 809 frames from 22 videos were extracted and used to train the network, achieving a final train error of 2.08 pixels and a test error of 2.70 pixels. Raw DLC trajectories were further processed to reduce tracking noise. Frames with likelihood values below 0.3 were excluded and temporarily assigned as missing values. Missing coordinates were subsequently interpolated linearly between adjacent valid frames. To remove isolated tracking spikes, a nearest-neighbor outlier correction algorithm was applied. Individual frames were identified as artifacts when neighboring frames remained spatially similar while the intermediate frame exhibited a large displacement. In these cases, coordinates were replaced by the average of the adjacent frames. To further constrain implausible tracking jumps, maximum frame-to-frame displacement was limited to 30 pixels per frame. Finally, trajectories were smoothed using a median filter with a kernel size of 5 frames to reduce high-frequency noise. Mouse speed was calculated from the frame-to-frame displacement of the body center and the maximum speed value was capped at 20 cm/s. Mouse angular velocity was calculated from vectors connected the body center to the nose and tail base. DLC network is available **(see Data and code availability)**.

### Immunohistochemistry and Microscopy

Mice were perfused with ice-cold 4% paraformaldehyde in phosphate-buffered saline. Serial horizontal or sagittal sections (50 µm) were cut on a sliding microtome (Leica SM2000R). Sections were first incubated in a blocking solution containing Normal Goat Serum (10%, Jackson) and TritonX-100 (0.05%, Sigma) for 2 h at room temperature. They were then incubated with primary antibodies overnight at 4°C. nRT cells were labeled with mouse anti-PV antibody (1:1000, Swant, PV27). KCC2 protein was stained by rabbit anti-KCC2 antibody (1:500, Sigma-Aldrich, 07-432). After wash, Sections were further incubated with secondary antibodies for 2 h at room temperature. Secondary antibodies used were goat anti-mouse-Alexa 594 (1:1000, Invitrogen, A11032) and goat anti-rabbit-Alexa 488, (1:1000, Invitrogen, A11008). Sections were mounted in an antifade medium (Vectashield) and images were obtained with Keyence BZ-9000 microscope. Image analyses were performed with ImageJ.

## Quantification

### *Ex vivo* Slice electrophysiology

*Characterization of minimal oPSCs*: To identify successfully evoked minimal oPSCs, we first applied a gaussian filter to smooth the raw traces, using a convolution kernel with a 5 ms standard deviation. Traces containing events that preceded the stimulus onset or exhibited unstable baselines were excluded. Successful events were identified in the smoothed traces using a dynamic threshold set at five times the standard deviation of the pre-stimulus baseline. Only these successful traces were used to compute the mean trace, using the raw (unsmoothed) data. From the mean trace, the following synaptic current features were extracted:

- Latency: The time from stimulus onset to 10% rise of the oPSC.
- Amplitude: The peak oPSC amplitude.
- Rise time: The time between 10% and 90% of the oPSC rise.
- Charge: The integral of the oPSC from 10% of the oPSC rise to the return to baseline.
- Decay tau(double-exponential, weighted): Obtained by fitting a double-exponential decay function to the oPSC waveform and calculating the weighted time constant:

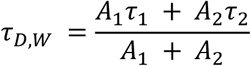

where A_1_ and A_2_ are the slow and fast amplitude components, and τ_1_ and τ_2_ are the slow and fast decay-time constants, respectively.

#### Detection of extracellular spikes in cell-attached configuration

To isolate negative-going voltage deflections corresponding to extracellular spikes, the voltage trace was inverted to convert troughs to peaks, and peaks were detected using the find_peaks function from the SciPy library. The peaks were detected based on a dynamic threshold set to 70% of the maximum data point in each sweep.

#### Detection of action potentials in whole-cell configuration

Action potentials were detected based on the first derivative of the voltage trace. Action potential onsets were identified as time points where dV/dt first exceeded 50 V/s.

### Fiber photometry

#### Preprocessing

The first 10 seconds of each recording were discarded to avoid onset artifacts. Signals were low-pass filtered (Butterworth, 2^nd^ order, 2 Hz cutoff) to reduce high-frequency noise, then fit with a double exponential function to capture and remove slow baseline drift. To correct for motion artifacts, we performed linear regression between the detrended 405 nm and 465 nm signals. The scaled 405 nm trace (motion estimate) was subtracted from the detrended 465 nm signal to yield a motion-corrected GCaMP trace. This corrected signal was then z-scored for normalization across sessions. To align fiber signals with movement, the corrected and z-scored GCaMP trace was temporally resampled to match the movement trace by computing the mean GCaMP signal within each video frame interval.

#### Detection of movement bouts

To identify discrete movement bouts, we applied a two-threshold algorithm. First, a low threshold of 2 cm/s was used to detect movement onset and offset. Onsets were defined as time points where speed rose above this threshold, and offsets as time points where speed subsequently fell below it. To ensure that only robust movements were captured, a secondary criterion required that the peak speed within each detected bout exceed 5 cm/s. Bouts not meeting this criterion were excluded from analysis.

#### Detection of left/right turns

To identify discrete turning events, angular velocity traces were thresholded at ±π/2 rad/s. Left turns were defined as timepoints in which angular velocity fell below −π/2 rad/s, whereas right turns were defined as periods in which angular velocity exceeded +π/2 rad/s. .

### Statistical analyses

General graphing and statistical analyses were performed using Python and Prism (GraphPad). Sample size is defined by the number of observations (i.e., cells or mice). No statistical method was used to predetermine sample size. Data are presented as mean ± standard error of means, unless specified otherwise. Normal distributions of data were not assumed. Animal subjects and cell recordings were randomized within experimental blocks to yield approximately equal sampling of experimental conditions. Statistical significance values and sample sizes are described in the figure legends. Statistical thresholds used were as follows: * *P* < 0.05; ** *P* < 0.01; *** *P* < 0.001; **** *P* < 0.0001; *n.s.* Not Significant.

## Acknowledgment

We are grateful to László Acsády and Nóra Hádinger for discussions on KCC2. We thank Massimo Scanziani, Alexandra Nelson, Vikaas Sohal, John R. Huguenard, C. Savio Chan, and members of the Paz Lab for feedback on the manuscript. We thank Reuben Thomas for statistical consultations and Misha Zilberter for technical advice on perforated-patch. We thank Aryn Gittis for providing the Npas1-iCre transgenic mouse lines. We thank Francoise Chanut and Kathryn Claiborn for editorial feedback. Funding sources: Croucher Foundation for Doctoral Studies (IYMC), Gladstone Institutes’ endowed funds (JTP).

## Author Contributions

IYMC and JTP conceived the study. IYMC performed all experiments and analyses. IYMC and JTP wrote the paper.

## Data and code availability

All data, code, and DLC network are available on https://zenodo.org/records/20583270 and https://github.com/isaachkchang/GPe-nRT.

## Figure legends

**Supp Fig 1.**
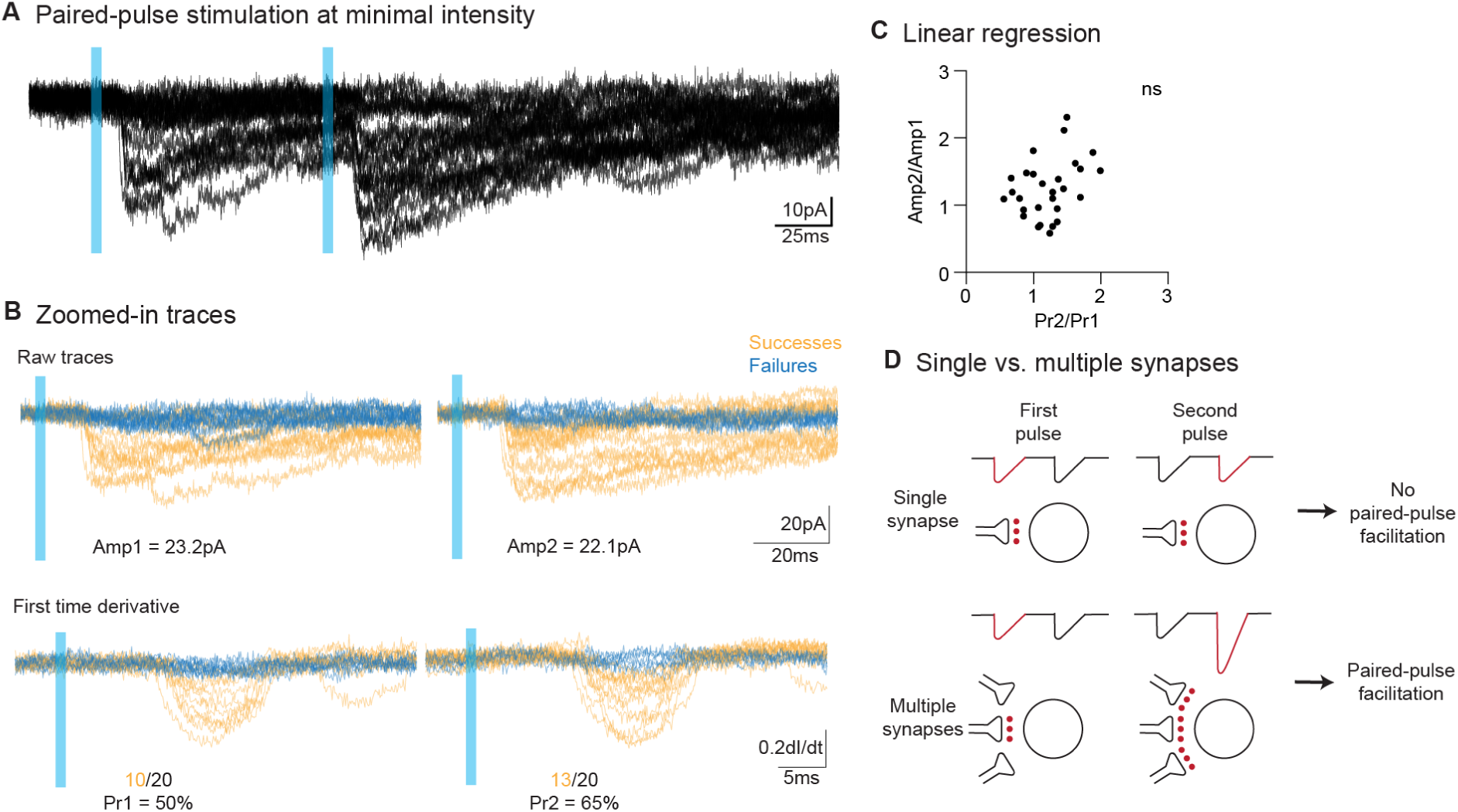
Minimal stimulation protocol reflects unitary transmission at GPe→nRT synapse. **A,** Representative traces of minimal oPSCs recorded from an nRT neuron in response to paired-pulse stimulation of a single GPe axon. **B,** Top, Expanded view of raw traces, colored by successes (yellow) and failures (blue). Bottom, Same as Top, but for the first time derivative. **C,** Scatter plot showing the correlation between the paired-pulse ratio (second oPSC amplitude/first oPSC amplitude) and paired-pulse facilitation (second release probability/first release probability). Linear regression (*n* = 29 cells, *P* = 0.11). **D,** Illustration of the principle of minimal stimulation. For two or more synapses, as release probability increases, both synapses tend to release quanta simultaneously, resulting in a higher amplitude for the second oPSC. However, for a single synapse, the amplitude of the second oPSC remains unchanged despite increasing release probability. For reference, see(*32*).

**Supp Fig 2.**
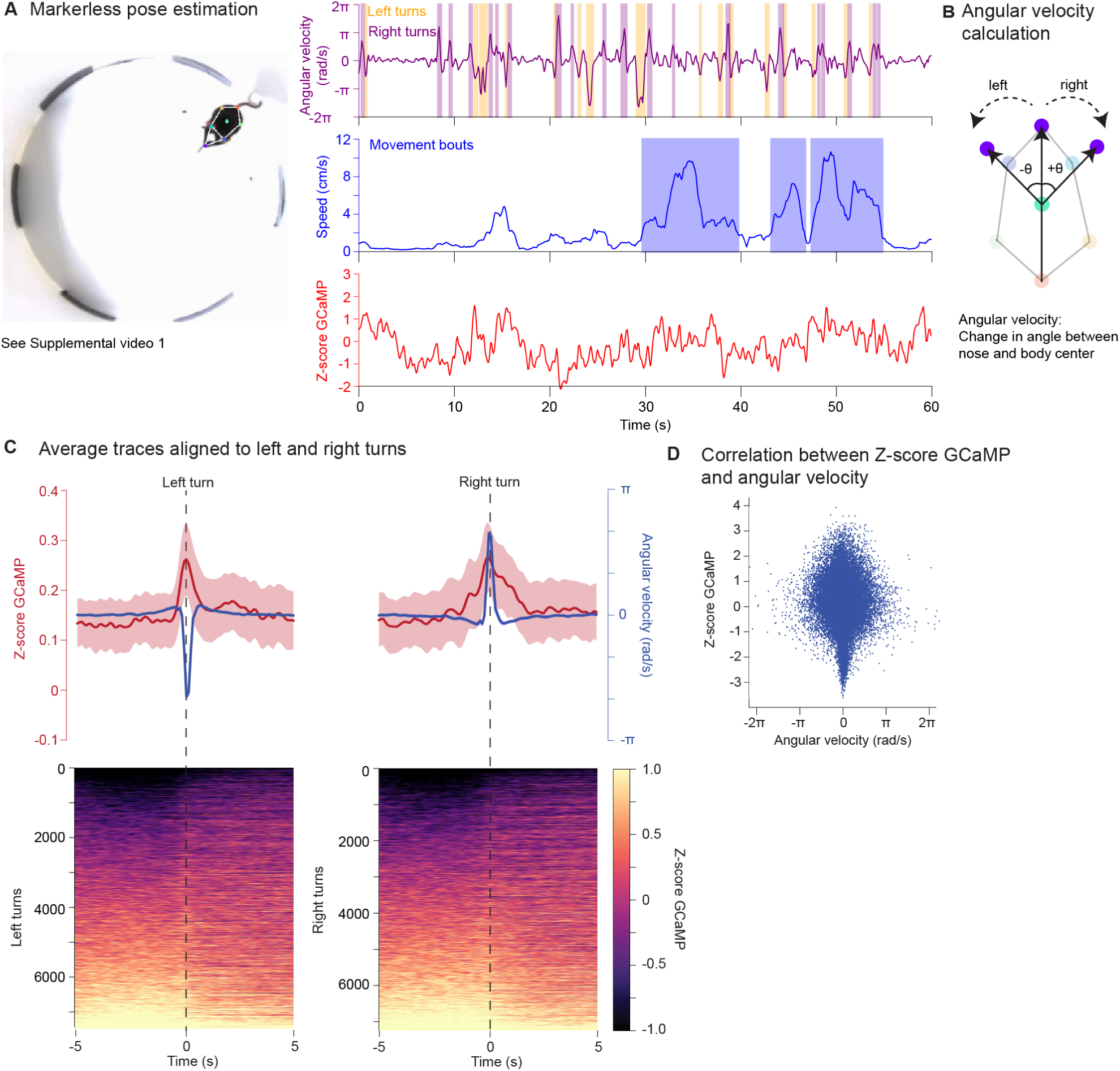
Turn-related Ca2+ activity at GPe→anterior nRT terminals. **A,** Video snapshot of a mouse in an open-field arena tethered to patch cords for fiber photometry. Color circles represent pose estimation markers. Right Simultaneous tracking of angular velocity, speed, and Ca^2+^ activity in GPe→anterior nRT terminals. **B,** Illustration of the calculation of angular velocity. **C,** Z-score GCaMP aligned with left turns (left, left, *n* = 7434 bouts, *N* = 10 mice) and right turns (right, *n* = 7225 bouts, *N* = 10 mice). Top, Mean traces ± SEM. Bottom, Heatmaps of Z-score GCaMP sorted based on pre-left/right-turn activity. **D,** Top, Scatter plot showing the relationship between Z-score GCaMP and angular velocity from one recording. Each marker is one frame (1/10 s).

## Tables

**Supp Table 1| Summary statistics**

## Videos

**Supp Video 1| Simultaneous tracking of GPe→nRT axonal activity, mouse speed and angular velocity**

